# Simultaneous transcriptome and proteome profiling in a single mouse oocyte with a deep single-cell multi-omics approach

**DOI:** 10.1101/2022.08.17.504335

**Authors:** Yi-Rong Jiang, Le Zhu, Lan-Rui Cao, Qiong Wu, Jian-Bo Chen, Yu Wang, Jie Wu, Tian-Yu Zhang, Zhao-Lun Wang, Zhi-Ying Guan, Qin-Qin Xu, Qian-Xi Fan, Shao-Wen Shi, Hui-Feng Wang, Jian-Zhang Pan, Xu-Dong Fu, Yong-Cheng Wang, Qun Fang

## Abstract

Nowadays, although single-cell multi-omics technologies are undergoing rapid development, simultaneous transcriptome and proteome analysis of a single-cell individual still faces great challenges. Here, we developed a single-cell simultaneous transcriptome and proteome (scSTAP) analysis platform based on microfluidics, high-throughput sequencing and mass spectrometry technology, to achieve deep and joint quantitative analysis of transcriptome and proteome at the single-cell level for the first time. This platform was applied to analyze single mouse oocytes at different meiotic maturation stages, reaching an average quantification depth of 19948 genes and 2663 protein groups in single mouse oocytes. This reliable quantitative two-omics dataset of single cells provided an important resource for understanding the relationship between the transcriptome and the proteome in cells. Based on the correlation analysis of RNAs and proteins in the same single cell, we demonstrated the expressive heterogeneity of transcriptome and proteome during the cellular biological process. Specially, we analyzed the meiosis regulatory network during oocyte maturation with an unprecedented depth at the single-cell level, and identified 30 transcript-protein pairs as specific oocyte maturational signatures, providing crucial insights into the regulatory features of transcription and translation during oocyte meiotic maturation.

## Introduction

Cellular heterogeneity is a fundamental property of various cellular systems, which not only refers to the genetic heterogeneity, but also includes the heterogeneity of cellular components during biological processes. Single-cell multi-omics technology can provide an effective tool to deeply recognize the cellular heterogeneity, benefiting the comprehensive understanding of the natural laws of human life activities^1–2^. Nowadays, various single-cell RNA sequencing techniques including 10× Genomics^3^, Smart-seq2^4^, and Seq-well^5–6^ have been developed and are used in studies of single-cell heterogeneity^7–8^ and cell atlas^9–10^. Meanwhile, via the combination of the single-cell RNA sequencing technique with the genome and epigenome sequencing technique, multi-omics sequencing of genome, epigenome, and transcriptome at the single-cell level has been achieved, such as the DR-seq^11^, G&T-seq^12^, scMT-seq^13^, scM&T-seq^14^, and sci-CAR^15^ techniques.

In spite of the significant progress of single-cell transcriptome and epigenome sequencing techniques, the single-cell proteomic analysis technique is still in the developing stage. Unlike nucleic acids, proteins are biochemical components that cannot be amplified, which presents great challenges to the analytical techniques, especially in the analysis of single-cell samples usually with a total protein content of around 200 pg in a mammalian somatic cell. At present, the targeting strategy using specific antibodies to label the targeted proteins is frequently adopted to achieve singlecell proteomic analysis, and several types of labeling antibodies are widely used, including fluorescent-labeled antibodies in imaging^16^, inorganic element-labeled antibodies in CyTOF^17^, and nucleic acid-labeled antibodies in high-throughput systems^18^. Most of these techniques can typically detect tens to hundreds of pre-targeted proteins in single cells, due to the limitation in the number of different antibody species and their specificity. The mass spectrometry (MS)-based technique coupled with the shotgun strategy provides an effective way to achieve non-targeted and deep single-cell proteome analysis. Recently, some MS-based systems such as SCoPE-MS^19^, Nested Nanowell chip^20,21^, Evosep^22^, and PiSPA^23^ could achieve much deeper proteomic analysis with over 1000 protein groups identified in single cells, and even reaching up to 3000 protein groups using the PiSPA platform. These progress demonstrated a significant breakthrough in single-cell protein identification depth, although they have not reached a comparable level to transcriptome sequencing.

According to the “central dogma”, mRNA is the template for protein synthesis, and proteins are the direct executor of cellular functions. Currently, it is still difficult to accurately predict the expression of proteins from transcriptional information alone. Therefore, simultaneous analysis of transcriptomes and proteomes in single cells is essential for the comprehensive characterization of cellular activities. However, despite the great progress in single-cell transcriptome and proteome analysis technology, simultaneous transcriptome and proteome analysis in single-cell individuals is still challenging, due to the extremely low amounts of total mRNAs and proteins in single cells as well as the difficulty in the cooperation between the two-omics analysis operations. Recently, a few techniques for simultaneous transcriptome and proteome analysis of single cells have been developed mainly combining the nucleic acid-labeled antibody-based protein identification approach with the RNA sequencing technique, such as CITE-seq^24^ and REAP-seq^25^, achieving simultaneous analysis of mRNAs and cell surface proteins in single cells. In addition, in order to analyze the intracellular proteins, the SCBC approach has been developed based on microfluidic chips^26^, in which the antibodies are immobilized in the chamber of the chip to capture and measure specific proteins in single-cell lysates, while free mRNAs are enriched and analyzed using the magnetic beads-based sequencing method. InCITE-seq^27^ is another type of intracellular protein analysis method developed on the basis of the CITE-seq technique, using modified double-chain nucleic acids binding antibodies for simultaneous analysis of proteins and transcriptome inside single cells. In addition, DBiT-seq^28^ and DSP^29^ are spatial multi-omics techniques for simultaneous transcriptome and proteome analysis of tissue sections at single-cell resolution from 10 μm to 50 μm. Since these multi-omics techniques are all developed based on antibody strategy for the analysis of proteins, the analysis depth of proteomics ranged from 4 to 82 targeted proteins, which is still limited in many biomedical researches.

In view of the advantages of the MS-based proteomic analysis technique in non-targeted and in-depth proteome analysis, the MS-based single-cell multi-omics technique has undergone rapid progress in recent years. Due to the incompatibility of principle and method between MS technique and RNA sequencing technique, current single-cell multi-omics studies based on MS technique usually conduct parallel analysis of transcriptome and proteome for different batch cells from the same samples. On the basis of the SCoPE-MS technique, Slavov’s group has developed the SCoPE2 technique^30^ and achieved the parallel analysis of single-cell transcriptome and proteome in differentiated monocyte samples with two batches of cells by combining the SCoPE2 technique with 10× Genomics. It is generally believed that the ideal single-cell multi-omics analysis should be capable of acquiring the multi-omics information from the same single-cell individual, rather than from different cells in the same sample. However, to achieve simultaneous multi-omics analysis in a single cell using the mass spectrometry technique based on the shotgun strategy still faces great technical challenges, owing to its complex multi-step sample pretreatment as well as the difficulty in cooperating with the sequencing operation.

Here, we developed a single-cell simultaneous transcriptome and proteome (scSTAP) profiling platform based on the microfluidic and label-free shotgun proteomic technique. This platform is capable of completing single cell capture, enzyme-assisted cell lysis, nanoliter-scale precise sample splitting, MATQ-seq^31^-based single-cell transcriptomic analysis and MS-based single-cell proteomic analysis. With the platform, we achieved deep and quantitative transcriptome and proteome analysis of single mouse oocytes. Based on the two-dimensional data obtained from each individual cell, the interaction networks of transcriptome and proteome during oocyte meiotic maturation were analyzed with an unprecedented depth.

## Results and discussion

### Simultaneous analysis of transcriptome and proteome in single cells

The aim of this work is to develop a platform and workflow to achieve the simultaneous analysis of MS-based proteomes and full-length sequencing-based transcriptomes in the same single-cell individuals. At present, single-cell proteome analysis technique plays a speed-determining role in the development of single-cell multi-omics analysis techniques, because the amount of proteins contained in a single cell is extremely small and proteins cannot be amplified, leading to greater challenges than single-cell transcriptome analysis. For the analysis of transcriptome and proteome in a single-cell individual, the primary problem to be solved is how to separate and transfer the very small amount of RNAs and proteins existing in the single cell for respective transcriptome sequencing and proteome analysis.

Based on our previously-developed sequential operation droplet array (SODA) technique^32,33^, which enables precise metering and manipulation of liquids in the nanoliter to picoliter range, we developed an approach called precise sample splitting (PSS) to achieve the quantitative division of single-cell samples in the nanoliter range for simultaneous transcriptome and proteome analysis. To realize such a straightforward multi-omics analysis strategy adopting the precise splitting of microsamples for separate multi-omics analysis, two prerequisites should be met. One prerequisite is that the system should have the ability to precisely split single-cell lysates in the nanoliter range, since nanoliter-scale sample volumes were usually adopted in most of the reported single-cell proteome analysis systems to depress the excessive dilution and the adsorption loss of ultra-trace proteins. Only when a system has such a quantitative sample splitting capability can it precisely and reproducibly control the splitting volumes and ratios for each single-cell samples, so as to ensure the comparability between the datasets of transcriptome and proteome obtained from different single cells. The other prerequisite is that after the single cell is lysed and before the lysate is divided, the cellular components should be uniformly distributed in the single-cell lysate solution. Otherwise, even if the sample splitting can be performed quantitatively, the uneven distribution of the single-cell components such as RNAs and proteins will cause inconsistency in the component content between the two aliquots. This will lead to inaccurate quantitative results, deteriorating the reliability of singlecell multi-omics analysis results and the comparability of datasets from different singlecell samples. Besides the two prerequisites, a higher requirement for the PSS-based single-cell multi-omics analysis is that the performance of the simultaneous two-omics analysis should not have a remarkable decrease in the identification depth compared with the single-omics analysis.

For the precise liquid manipulation in the single-cell multi-omics analysis, we used an improved SODA system to perform the precise sample splitting and multi-step sample pretreatment. Its precision in manipulating hundreds of nanoliter of liquids (e.g. 500 nL) was 5.4% (coefficient of variation [CV], n = 12), which could meet the requirement for quantitative splitting of single-cell samples in the present work. For the choice of the reactor, in order to reduce the loss of RNAs and proteins absorbed on the surface of the reactors, as well as the loss during the multi-step pretreatment process, we choose the commercial insert tubes with a tapered bottom and hydrophobic surface in each tube as the stationary droplet reactors, which could be directly coupled with the commercial autosampler of the liquid chromatography. The tapered bottom of the insert tubes was used to load the nanoliter-scale sample droplets and conduct in-situ multistep sample pretreatment.

### Optimization and Performance of the scSTAP platform

The workflow of scSTAP platform is shown in Figure 1, including the steps of single cell capture, cell lysis and sample splitting, transcriptome analysis, and proteome analysis. In the cell lysis, good results were obtained in the consistency analysis of both proteome and transcriptome in two splitting droplets with enzyme assisted lysis (Figure 2A, Figure S1), which validated the reliability of the quantification results of proteome and transcriptome as well as the applicability of this scSTAP platform in the joint analysis of single-cell multi-omics. In addition, we compared the quantitative results between an intact and a half of cells. In order to minimize the effect of different cell states on the experimental results, we chose the fully-grown oocytes at a stably biological state as the tested samples. The results showed that there was no significant difference between the identification depth of the intact and the half of cells (Figure 3A, Figure S2).

**Fig. 1.**
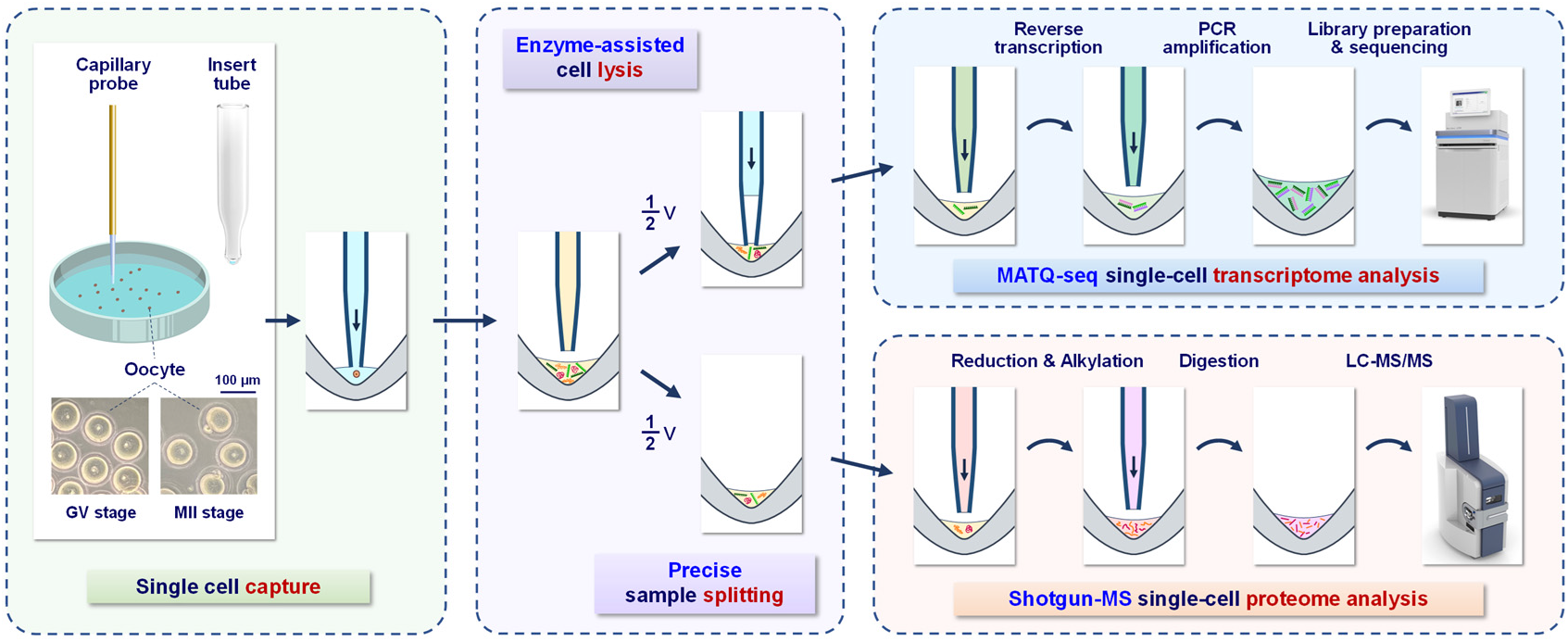
Schematic diagram of the scSTAP platform and workflow for single-cell multi-omics analysis. The workflow includes the steps of single cell capture, cell lysis and sample splitting, transcriptome analysis, and proteome analysis. In the single cell capture, a capillary probe of the droplet manipulation module was used to pick up a single target cell into an insert tube reactor. In the cell lysis, the Lys-C and RapiGest solutions were used to lyse the single cell sample. In the precise sample splitting, the capillary probe was used to precisely split the cell lysate into two aliquots. The transcriptome of one aliquot was analyzed by the MATQ-seq workflow including the reverse transcription, PCR amplification, library preparation, and DNA sequencing. The proteome of the other aliquot was analyzed using shotgun proteomics method including protein reduction, alkylation, digestion and LC-MS/MS analysis.

**Fig. 2.**
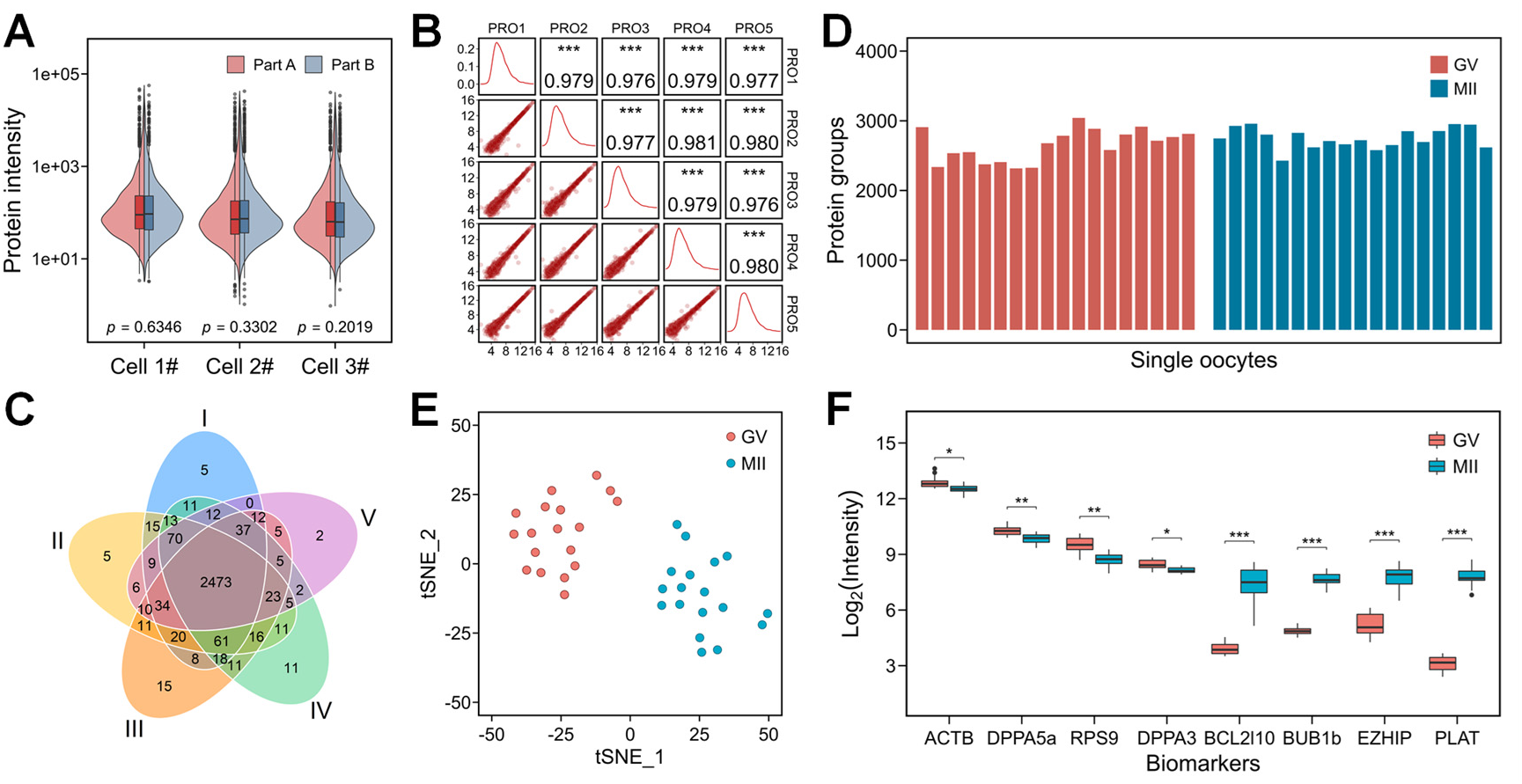
Performance of proteome analysis of the scSTAP platform. (A) Consistency analysis of single-cell sample splitting in proteome analysis. (B) Correlation coefficient of proteome expressions in the QC sample (n = 5). (C) Venn diagram of the identified protein groups in the QC sample (n = 5). (D) Identification numbers of protein groups in the oocyte samples at GV stage (n = 18) and MII stage (n = 18). (E) tSNE cluster visualization in single-cell proteome analysis of the oocyte samples at GV stage (n = 18) and MII stage (n = 18). (F) Comparison of the typical protein intensity in the oocyte samples at GV stage (n = 18) and MII stage (n = 18). ***, *p* = 6.08 × 10^−27^ for PLAT, *p* = 3.67 × 10^−21^ for BUB1b, *p* = 1.38 × 10^−14^ for EZHIP and *p* = 7.90 × 10^−13^ for BCL2l10. **, *p* = 2.17 × 10^−3^ for RPS9 and *p* = 2.12 × 10^−3^ for DPPA5a. *, *p* = 1.51 × 10^−2^ for DPPA3 and *p* = 1.05 × 10^−2^ for ACTB.

**Fig. 3.**
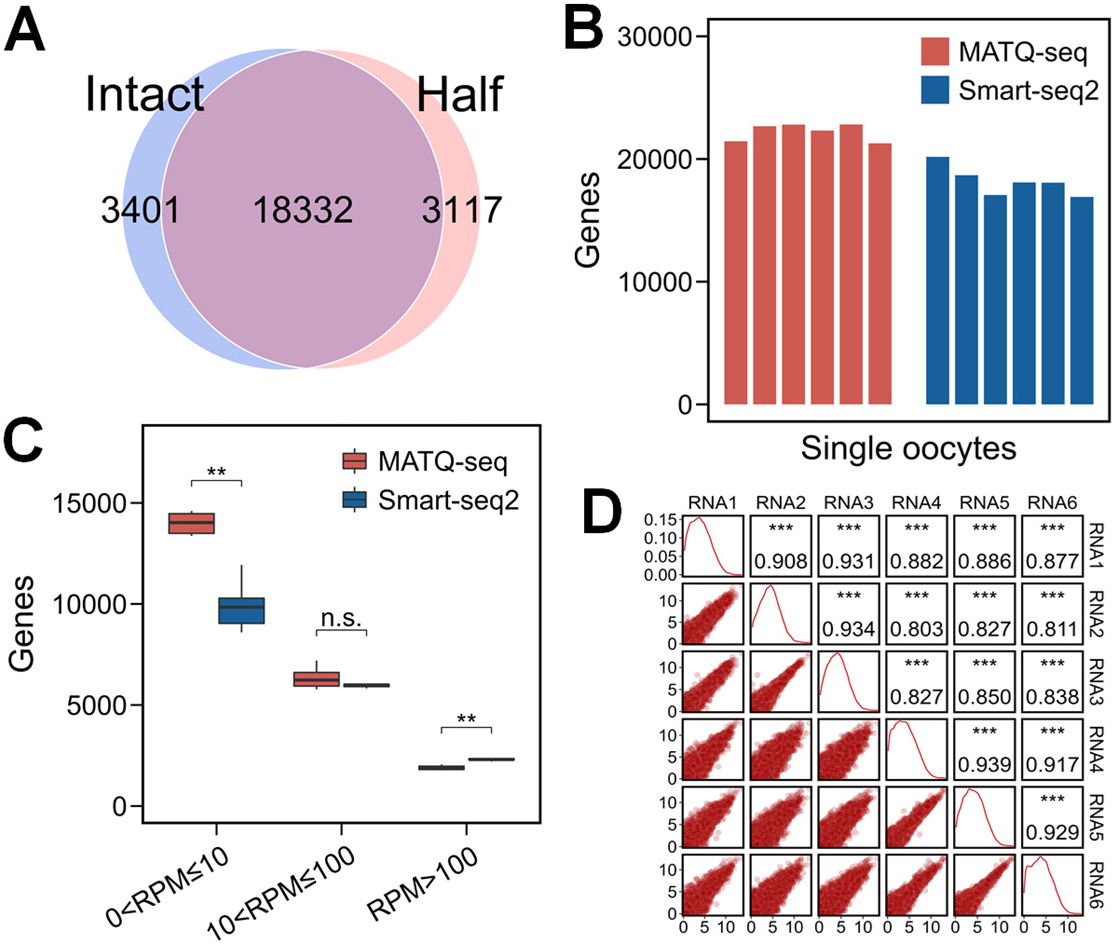
Performance of transcriptome analysis of the scSTAP platform. (A) Venn diagram of the identified genes in the intact single oocyte and a half of single oocyte samples. (B) Identification numbers of genes in single oocyte samples at GV stage using MATQ-seq (n = 6) and Smart-seq2 (n = 6). (C) Comparison of the gene numbers using MATQ-seq (n = 6) and Smart-seq2 (n = 6) in single oocyte samples at GV stage. **, *p* = 0.0022 for 0 < RPM ≤ 10, and *p* = 0.0022 for RPM > 100. Not significant (n.s.), *p* = 0.31 for 10 < RPM ≤ 100. (D) Correlation analysis of gene expressions in the single oocyte samples using MATQ-seq (n = 6).

With the optimized conditions, the data independent acquisition (DIA) mode was used for single oocyte proteomic analysis. For validation of this workflow, we prepared a quality control (QC) sample containing forty mouse oocytes and analyzed it repeatedly with an injection amount equivalent to a single oocyte for each run. In the correlation analysis of the QC sample (n = 5), the correlation coefficients (CCs) of proteome expressions in the QC sample were all higher than 0.97 (Figure 2B). According to the results of Venn diagram, 85% of the total protein groups had been quantified in all runs (Figure 2C). In addition, the median variation coefficient of these quantified protein groups was less than 20% among five replicates (Figure S3). These results indicated that this scSTAP platform had a good repeatability in proteome analysis.

To evaluate the performance of this scSTAP platform in actual proteome analysis, we applied it to analyze single mouse oocytes at different maturation stages, including a total of 18 oocytes at fully grown germinal vesicle (GV) stage and 18 oocytes arrested in second meiotic metaphase (MII) stage. In the proteomics analysis, we quantified an average of 2703 proteins in all the single oocyte samples (Figure 2D). After the data accumulation, a total of 3363 proteins were quantified in these oocyte samples, among which 1600 proteins expressed in all the GV samples and 1927 proteins expressed in all the MII samples. Based on the expression matrix of the quantified proteins, oocytes at GV and MII stages could be well clustered using the unsupervised cluster approach, and the clustering result was consistent with the morphological analysis result (Figure 2E). Similar to the reported results^34^, some stable proteins in oocytes, such as ACTB, DPPA5a, RPS9, and DPPA3 had low variations of expression (CV < 30%) among all the single-cell samples, while the expression of some variable proteins had a significant difference between oocytes at GV and MII stages (Figure 2F), demonstrating the reliability of the proteome quantification data obtained by the scSTAP platform.

The performance of MATQ-seq used in the present workflow was evaluated by comparing its sequencing results with that using the Smart-seq2 method. In the identification performance, an average of 22224 and 18168 genes were identified in single oocyte samples with the MATQ-seq (n = 6) and Smart-seq2 (n = 6) methods, respectively (Figure 3B). Especially, for the low-abundance genes with the reads per million mapped reads (RPM) less than 10, the performance of the MATQ-seq method was significantly higher than that of the Smart-seq2 method (Figure 3C). In addition, there was no obvious 3’- or 5’-end bias observed in the mapped read coverage of the MATQ-seq method (Figure S4). In the quantification performance, the CCs of RPM in the single oocyte samples (n = 6) were all higher than 0.80 (Figure 3D).

### Deep transcriptome and proteome profiling of single mouse oocytes

With the optimized scSTAP platform and workflow, we conducted the single-cell multi-omics analysis to achieve the simultaneous deep transcriptome and proteome profiling of mouse oocytes at the GV and MII stages. An average quantification depth of 19948 genes and 2663 protein groups were obtained in single oocytes (Figure 4A). To explore the variance of oocytes during meiotic maturation, the single-cell multi-omics profiles were subjected to clustering analysis. In the principle component analysis (PCA) of the transcriptome dataset, the main contributing genes of the first principal component (PC1) were DCAF1 for positive and PRDX2 for negative. Correspondingly, the top contributing genes of the first principal component (PC1) in the PCA of the proteome dataset were CPEB1 and BPGM. However, based on these PCA results for transcriptome (Figure 4B) and proteome (Figure 4C), the shared nearest neighbors (SNN) algorithm could not distinguish the single oocytes at GV and MII stages significantly, and the information from transcriptome and proteome could not be jointly analyzed. To further conduct the joint analysis of the multi-omics dataset, we analyzed the transcriptome and proteome expression matrix of single mouse oocytes based on the unsupervised weighted nearest neighbor (WNN) clustering analysis^35^. Based on the matrix of weighted coefficient for the transcriptome and proteome, the contribution of both the transcripts and proteins could be well taken into account in the joint analysis of multi-omics. The UMAP visualization results showed that single oocyte samples at GV and MII stages could be precisely distinguished using the WNN algorithm (Figure 4D).

**Fig. 4.**
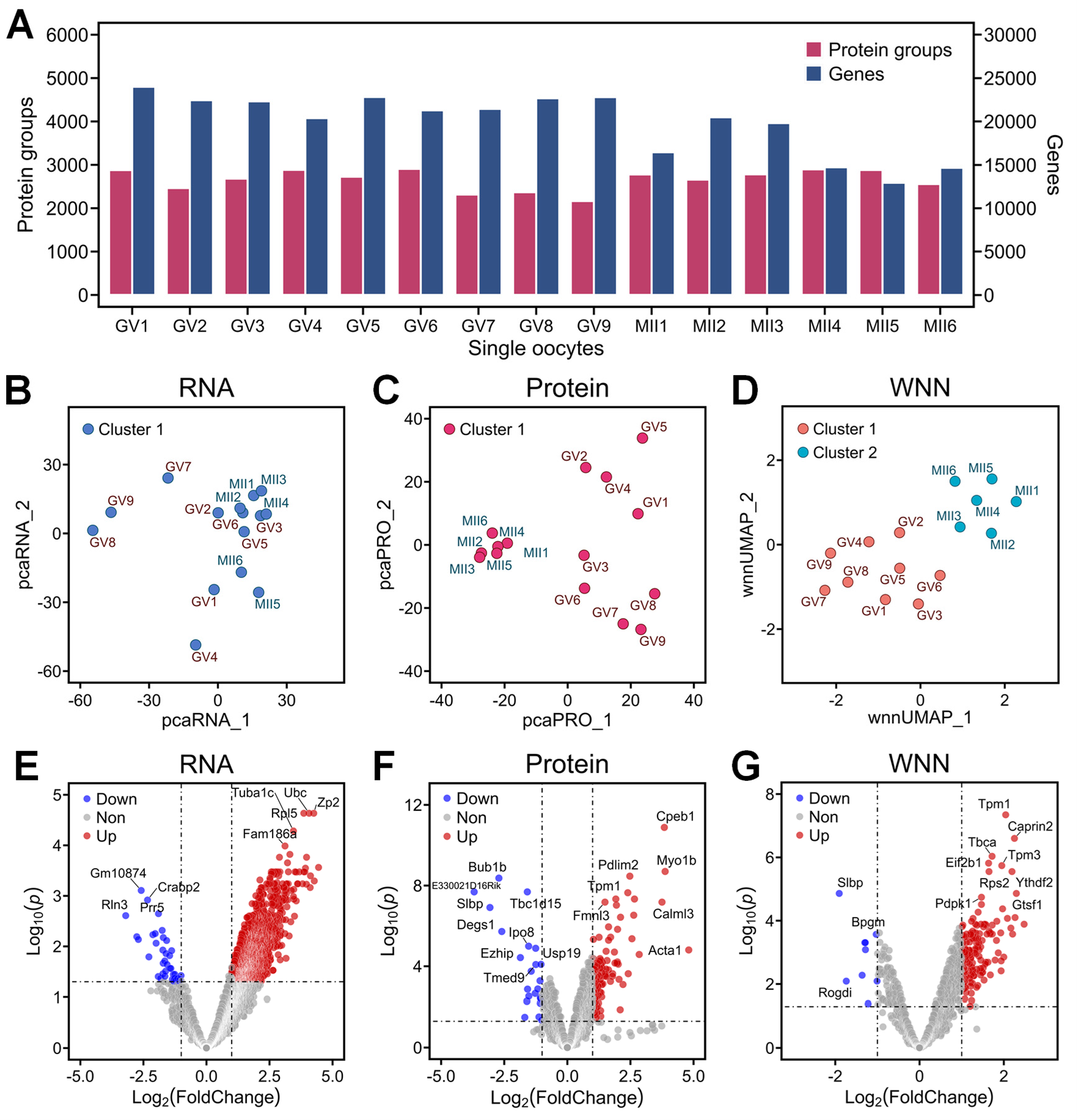
Cluster analysis and differential expression analysis of single mouse oocytes during meiotic maturation. (A) Identification numbers of protein groups and genes in single mouse oocytes at GV stage (n = 9) and MII stage (n = 6). In average, 2599 ± 278 protein groups and 22242 ± 1059 genes were quantified in the single mouse oocytes at GV stage. In addition, 2759 ± 130 protein groups and 16509 ± 3037 genes were quantified in the single mouse oocytes at MII stage. (B-C) Principal component analysis of single mouse oocytes at GV and MII stages based on expression matrix of transcriptome (B) and proteome (C). (D) UMAP cluster analysis of single mouse oocytes at GV and MII stages for the multi-omics joint analysis based on the weighted nearest neighbor (WNN) algorithm. (E-F) Volcano diagrams of differential expressed transcripts (E) and proteins (F) in single mouse oocytes at GV and MII stages. (G) Volcano diagrams of differential expressed genes in both transcriptome and proteome from single mouse oocytes during meiotic maturation.

To further investigate the dynamic patterns of multi-omics profiles, we analyzed the differentially expressed transcripts and proteins in single oocytes at GV and MII stages. Based on the statistical calculation, 2392 differentially expressed transcripts (Figure 4E) and 127 differentially expressed proteins (Figure 4F) were identified between GV and MII oocytes using the limma package in R (*p* < 0.05; fold change [FC] of log2 expression > 2). Particularly, based on the matrix of weighted coefficient from the WNN algorithm, we obtained the joint expression matrix of transcriptome and proteome. On the basis of this joint expression matrix, 166 differentially expressed RNA-protein pairs were identified (Figure 4G), and 30 transcript-protein pairs were identified as specific oocyte maturational signatures. Among these pairs, 5 related genes were known biomarkers including CPEB1^36^, GTSF1^37^, GDF9^38^, CELF1^39^, and ZAR1l^40^, while other genes could be considered as potential biomarker candidates.

### Regulatory networks of single mouse oocytes during meiotic maturation

To understand the characteristics of multi-omics profiles in regulatory network analysis, we focused on the key components of the “Oocyte Meiosis” pathway under two dimensions involving the expressions of RNAs and proteins during oocyte meiotic maturation. In the protein-protein interaction analysis (Figure S5), the differential expression of the proteome was more significant than the transcriptome. Notably, CPEB1 and GTSF1 were significantly downregulated in both transcriptome and proteome, while the expression patterns of BUB1b, SLBP, H1foo, and RBX1 were downregulated at the RNA level and upregulated at the protein level. These results showed that the expression of RNAs and proteins were not always consistent in the cellular biological processes, and the expression level of proteins could represent the state of the cells much more realistically. In addition, the expression of FBXO43, MYT1, CDC20, and CDC25c were absent in the proteomic analysis results, which implied that the depth of the single-cell proteome analysis should be further improved.

Long non-coding RNAs (lncRNAs) were also identified in this single-cell multi-omics profiles, which could not be translated into proteins, but had important functions in regulatory. Based on the GENCODE database, an average of 3389 lncRNAs were expressed in the GV-stage oocytes and an average of 2568 lncRNAs were expressed in the MII-stage oocytes. According to the differential expression analysis results of the transcriptome, 21 differentially expressed transcripts were identified as lncRNAs and only the function of Platr14 had been reported to be associated with embryonic development^41^. Among these lncRNAs, the expression of C86187 and Gm1965 were high (RPM > 500) enough to merit further investigation.

## Conclusions

Single-cell multi-omics technique is an important means to explore life activities at the single-cell level. Current multi-omics approaches can only achieve the analysis of the genomes, epigenomes, and transcriptomes from the same single cells. In the present work, we developed the scSTAP-based single-cell multi-omics analysis platform to realize the simultaneous in-depth analysis of the transcriptome and proteome in single cells. The deep and quantitative transcriptome and proteome datasets from the same single-cell individuals were obtained for the first time. These data provide an unprecedented opportunity to understand the correlation between the transcriptome and proteome expression at the same individual level of single cells. Different from the conclusion obtained in previous studies that the transcriptome and proteome expression were poorly correlated, we found that even though the transcripts and proteins expressed of many (~50%) genes had low correlations (CCs < 0.3), at the individual gene level, the transcript and protein expression of some genes were highly correlated, and even the transcript expression of several genes were negatively correlated with the protein expression. These findings show that the transcription and translation process in the single-cell level is a complex multifactorial regulatory process, which provides crucial insights into the studies of cellular regulatory networks. We believe such a single-cell transcriptome-proteome analysis approach will provide a powerful tool with broad application in single-cell and biomedical research and will promote new breakthroughs in cellular and molecular biology, e.g. providing the solid data foundation for the study of the transcriptome-protein relationship—a fundamental biological question related to the central dogma.

Oocytes, as an important type of single-cell samples with reproductive and genetic ability, are one of the important objects in single-cell multi-omics research. At present, few research has been done on the proteome analysis of oocytes, and the analysis depths are still limited to hundreds of proteins^42,43^, while no research has been conducted on the simultaneous analysis of transcriptome and proteome in single oocytes. Our approach not only realized the simultaneous analysis of transcriptome and proteome in the same single mouse oocytes, but its analysis depth also exceeds the results of the conventional single-omics approaches^34,44^, which could provide important technical support for research on the reproductive development and quality assessment of oocytes.

In the future, the scSTAP approach could be further improved by adopting the multi-labeling transcriptome and proteome analysis techniques, such as using barcodes in RNA-sequencing and tandem mass tags (TMT) in proteomic analysis, to significantly increase the throughput of single-cell multi-omics analysis. Its application could also be extended to more different sources and types of single-cell samples. Furthermore, the present platform could also be further developed to achieve the simultaneous analysis of three-omics or more within a single-cell individual, such as genomics, epigenomics, transcriptomics, proteomics, and metabolomics analysis, by taking the advantage of the SODA technique in flexibly manipulating trace amounts of liquids as well as utilizing other existing single-cell omics analysis techniques.

## Supporting information

Supplementary information for scSTAP analysis

## Methods

### Establishment of the scSTAP platform

The scSTAP platform was composed by three modules, including a single-cell capture & droplet manipulation module, a single-cell transcriptomic analysis module and a single-cell proteomic analysis module. The single-cell capture & droplet manipulation module was built on the basis of the sequential operation droplet array (SODA) strategy previously developed in the authors’ group^32,33^. It consisted of a syringe pump, a capillary probe with a tapered tip and a microscope, which were installed on *x-y-z* translation stages. The single-cell transcriptomics analysis module included a thermocycler, a sonicator, and a sequencing system. The single-cell proteomic analysis module was composed of a nanoflow-liquid chromatograph with an autosampler and a trapped ion mobility spectrometry-mass spectrometer. A capillary column with an integrated MS spray tip was prepared with C_18_ stationary phase.

### Mice and collection of oocytes

The study followed the ethical guidelines of the Animal Research Committee of Zhejiang University. Wild-type (WT) female C57BL/6J mice were purchased from Beijing Weitahe Laboratory Animal Technology Co. All mice acclimated for a week in a controlled environment with 12 h light per day, air humidity of 50–70%, and temperature between 20 °C and 22 °C. To collect the oocytes at fully grown GV stage, around six-weeks-old to eight-weeks-old female mice were injected with 7 IU of pregnant mare serum gonadotropin (PMSG) and humanely sacrificed 44 h later. Then the oocytes at fully grown GV stage were collected in the M2 medium. For collecting the oocytes arrested in second meiotic metaphase (MII stage), approximately 48 h after PMSG (7 IU) injection, 7 IU of human chorionic gonadotropin was injected into female mice. After an additional 20 h, cumulus-oocyte complexes (COCs) were surgically removed from fallopian tubes. Then the oocytes at MII stage were collected in the M2 medium from COCs after digestion of 300 IU/mL hyaluronidase.

### Procedures of the scSTAP platform

A schematic workflow of the scSTAP platform for single-cell multi-omics analysis is shown in Figure 1. First, single oocyte cells were sucked into the capillary probe and pushed into the insert tube reactors to form nanoliter-scale droplets encapsulating a single cell in each droplet. Next, an enzyme assisted method was used to perform the cell pre-lysis by sequentially adding Lys-C and RapiGest solutions into each droplet. After the cell lysis, the droplets containing cell lyses were split into two aliquots using the capillary probe connected with the syringe pump. One aliquot was added into RNase inhibitor and Triton X-100 solutions for analysis of transcriptome, and the other aliquot left in the reactor was used for proteomic analysis.

In the single-cell transcriptomic analysis, the droplet was analyzed by the MATQ-seq technique^31^ including the reaction of reverse transcription and second-strand synthesis in PCR tubes. Briefly, in the reverse transcription, the reverse transcription mix was added into each droplet, and the droplets were treated with ten cycles of annealing (ramping from 8 °C to 50 °C) on the thermocycler. After reverse transcription, the residual primers were digested by T4 polymerase at 37 °C for 40 min and 80 °C for 20 min, and then the RNAs were digested by RNase-H and RNase-If at 37 °C for 15 min and 72 °C for 15 min. Next, dC-tailing and second strand synthesis were performed according to the protocol described in the MATQ-seq technique^31^. The PCR amplification of the droplets were conducted with a 28-cycle PCR program.

In the single-cell proteomic analysis, the droplet was analyzed using an improved deep single-cell proteomic technique developed by the authors’ group. Briefly, tris(2-carboxyethyl)-phosphine (TCEP), iodoacetamide (IAA), and trypsin solutions were sequentially added into the droplet to achieve protein reduction, alkylation, and enzymatic digestion.

### Library preparation and sequencing

After PCR amplification, pooled libraries were purified with 1.2X AMPure XP beads into PCR-grade water. The ABclonal Rapid Plus DNA Lib Prep kit of Illumina was used to construct the PCR product library according to the protocol provided with the kit. The cDNA was firstly sheared to 300 bp using a Covaris S220 sonicator. End repair and A-tailed was then performed using a NGS Fast DNA Library Prep kit of Illumina. T4 DNA ligase was used to ligate adaptors to samples. Libraries were then diluted and sequenced on a NovaSeq 6000 machine by Haprolos, using the NovaSeq S4 reagent kit v1.5 (300 cycles).

### LC-MS/MS analysis

After the pretreatment, the droplet containing the digested peptides was injected into the capillary column by the autosampler of the liquid chromatograph. The MS/MS analysis of the separated peptides was performed by the mass spectrometer equipped with a nano-spray ion source. The acquisition mode of diaPASEF was used in the analysis of the oocyte samples, with an isolation width of 25 m/z, and collision energy range of 20–47.3 eV in collision-induced dissociation (CID). For the conditions of data dependent acquisition (DDA) in the PASEF mode, the mass range was 300–1500 m/z and the capture width was 2–3 m/z, while the collision energy range was the same as the diaPASEF mode.

### Data analysis

The raw transcriptome sequencing data were trimmed using Cutadapt (version 4.0) to remove primer sequences, followed by trimming of the extra bases generated by dC-tailing. The reads were then mapped to the genome using STAR software (version 2.7.10a). Gene annotations were performed using Gene Code Annotation Release M29 (GRCm39, GENCODE). After retrieving the mapping information of the reads, feature counts were used to count the gene expression level. The numbers of total reads were summarized to normalize the gene expression. The unit of RPM is defined as the reads per one million total reads. Gene expression level is defined by the number of reads of a gene divided by the total number of reads of all genes and multiplied by 1,000,000. In addition, the raw data of the Smart-seq2 method were provided by LC-Bio Co. (Hangzhou, China).

The raw data of proteomics were analyzed by Spectronaut software (version 15.6.211220.50606) with the default settings against the UniProt database (UP000000589_10090.fasta, Mus musculus: 21,985 entries). The post analysis and visualization of bioinformatics data were conducted by corresponding R packages. The interaction networks were analyzed by STRING database (version 11.5) and visualized by Cytoscape software (version 3.9.1).

## Acknowledgements

The authors acknowledge the financial support provided by the National Natural Science Foundation of China (Grants 21827806 and 21435004), the National Ministry of Science and Technology (Grant 2021YFA1301601), the Leading Innovative and Entrepreneur Team Introduction Program of Zhejiang (Grant 2021R01012), the National Key Research and Development Program of China (Grant 2021YFC2700101), and the hundred talents program of Zhejiang University.

## Author contributions

Q.F., Y.C.W., X.D.F., and Y.R.J. conceived the idea. Q.W., J.B.C., Y.W., and H.F.W. built the platform. L.R.C. and X.D.F. collected the oocyte samples. Y.R.J., Q.W., and Z.Y.G. explored and optimized the LC-MS/MS conditions. Y.R.J., L.Z., L.R.C., Q.W., and F.Q.X. planned and performed experiments. Y.R.J., Q.W., Q.Q.X., and S.W.S. provided nano-LC columns. Q.F., Y.C.W., X.D.F., J.Z.P., and H.F.W. guided the research experiments and participated in the solution of problems arose in the experiments. Y.R.J., Q.W., L.Z., L.R.C., J.W., T.Y.Z., Z.L.W., Y.C.W., X.D.F., and Q.F. contributed to data analysis and interpretation, as well as figure preparation. Y.R.J., Q.F., L.Z., L.R.C., J.B.C., Y.C.W., and X.D.F. wrote and revised the manuscript. Q.F., Y.C.W., X.D.F., and J.Z.P. supervised the studies and acquired funding.

## Notes

### Competing Interest Statement

The authors have declared no competing interest.

## References

1. D. Mahdessian, A. J. Cesnik, C. Gnann, et al., Spatiotemporal dissection of the cell cycle with single-cell proteogenomics. Nature, 2021, 590, 649–654.

2. K. Grosselin, A. Durand, J. Marsolier, et al., High-throughput single-cell ChIP-seq identifies heterogeneity of chromatin states in breast cancer. Nat. Genet., 2019, 51, 1060–1066.

3. G. Zheng, B. Lau, M. Schnall-Levin, et al. Haplotyping germline and cancer genomes with high-throughput linked-read sequencing. Nat. Biotechnol., 2016, 34, 303–311.

4. S. Picelli, Å. Björklund, O. Faridani, et al., Smart-seq2 for sensitive full-length transcriptome profiling in single cells. Nat. Methods, 2013, 10, 1096–1098.

5. T. Gierahn, M. Wadsworth, T. Hughes, et al., Seq-Well: portable, low-cost RNA sequencing of single cells at high throughput. Nat. Methods, 2017, 14, 395–398.

6. X. Han, R. Wang, Y. Zhou, et al., Mapping the mouse cell atlas by microwell-seq. Cell, 2018, 172, 1091–1107.e17.

7. E. Papalexi, R. Satija, Single-cell RNA sequencing to explore immune cell heterogeneity. Nat. Rev. Immunol., 2018, 18, 35–45.

8. D. A. Lawson, K. Kessenbrock, R. T. Davis, et al., Tumour heterogeneity and metastasis at single-cell resolution. Nat. Cell Biol., 2018, 20, 1349–1360.

9. S. Petropoulos, D. Edsgärd, B. Reinius, et al., Single-cell RNA-seq reveals lineage and X chromosome dynamics in human preimplantation embryos. Cell, 2016, 165, 1012–1026.

10. A. Villani, R. Satija, G. Reynolds, et al., Single-cell RNA-seq reveals new types of human blood dendritic cells, monocytes, and progenitors. Science, 2017, 356, eaah4573.

11. S. Dey, L. Kester, B. Spanjaard, et al., Integrated genome and transcriptome sequencing of the same cell. Nat. Biotechnol., 2015, 33, 285–289.

12. I. Macaulay, W. Haerty, P. Kumar, et al., G&T-seq: parallel sequencing of singlecell genomes and transcriptomes. Nat. Methods, 2015, 12, 519–522.

13. Y. Hu, K. Huang, Q. An, et al., Simultaneous profiling of transcriptome and DNA methylome from a single cell. Genome Biol., 2016, 17, 88.

14. C. Angermueller, S. Clark, H. Lee, et al., Parallel single-cell sequencing links transcriptional and epigenetic heterogeneity. Nat. Methods, 2016, 13, 229–232.

15. J. Cao, D. A. Cusanovich, V. Ramani, et al., Joint profiling of chromatin accessibility and gene expression in thousands of single cells. Science, 2018, 361, 1380–1385.

16. M. Mondal, R. Liao, L. Xiao, et al., Highly multiplexed single-cell in situ protein analysis with cleavable fluorescent antibodies. Angew. Chem. Int. Edit., 2017, 56, 2636–2639.

17. S. C. Bendall, E. F. Simonds, P. Qiu, et al., Single-cell mass cytometry of differential immune and drug responses across a human hematopoietic continuum. Science, 2011, 332, 687–696.

18. A. S. Genshaft, S. Li, C. J. Gallant, et al., Multiplexed, targeted profiling of singlecell proteomes and transcriptomes in a single reaction. Genome Biol., 2016, 17, 188.

19. B. Budnik, E. Levy, G. Harmange, et al., SCoPE-MS: mass spectrometry of single mammalian cells quantifies proteome heterogeneity during cell differentiation. Genome Biol., 2018, 19, 161.

20. J. Woo, S. M. Williams, L. M. Markillie, et al., High-throughput and high-efficiency sample preparation for single-cell proteomics using a nested nanowell chip. Nat. Commun., 2021, 12, 6246.

21. Y. Cong, K. Motamedchaboki, S. A. Misal, et al., Ultrasensitive single-cell proteomics workflow identifies >1000 protein groups per mammalian cell. Chem. Sci., 2021, 12, 1001–1006.

22. A. D. Brunner, M. Thielert, C. Vasilopoulou, et al., Ultra-high sensitivity mass spectrometry quantifies single-cell proteome changes upon perturbation. Mol. Syst. Biol., 2022, 18, e10798.

23. Y. Wang, Z. Guan, S. Shi, et al., Pick-up single-cell proteomic analysis for quantifying up to 3000 proteins in a tumor cell. BioRxiv, 2022, doi: https://doi.org/10.1101/2022.06.28.498038.

24. M. Stoeckius, C. Hafemeister, W. Stephenson, et al., Simultaneous epitope and transcriptome measurement in single cells. Nat. Methods, 2017, 14, 865–868.

25. V Peterson, K. Zhang, N. Kumar, et al., Multiplexed quantification of proteins and transcripts in single cells. Nat. Biotechnol., 2017, 35, 936–939.

26. A. X. Xu, Q. Liu, K. L. Takata, et al., Integrated measurement of intracellular proteins and transcripts in single cells. Lab Chip, 2018, 18, 3251–3262.

27. H. Chung, C. N. Parkhurst, E. M. Magee, et al., Joint single-cell measurements of nuclear proteins and RNA in vivo. Nat. Methods, 2021, 18, 1204–1212.

28. Y. Liu, M. Yang, Y. Deng, et al., High-spatial-resolution multi-omics sequencing via deterministic barcoding in tissue. Cell, 2020, 183, 1665–1681.

29. C. R. Merritt, G. T. Ong, S. E. Church, et al., Multiplex digital spatial profiling of proteins and RNA in fixed tissue. Nat. Biotechnol., 2020, 38, 586–599.

30. H. Specht, E. Emmott, A. A. Petelski, et al., Single-cell proteomic and transcriptomic analysis of macrophage heterogeneity using SCoPE2. Genome Biol., 2021, 22, 50.

31. K. Sheng, W. Cao, Y. Niu, et al. Effective detection of variation in single-cell transcriptomes using MATQ-seq. Nat. Methods, 2017, 14, 267–270.

32. Y. Zhu, Y. X. Zhang, L. F. Cai, et al., Sequential operation droplet array: an automated microfluidic platform for picoliter-scale liquid handling, analysis, and screening. Anal. Chem., 2013, 85, 6723–6731.

33. Z. Dong, Q. Fang, Automated, flexible and versatile manipulation of nanoliter-to-picoliter droplets based on sequential operation droplet array technique. Trends Anal. Chem., 2020, 124, 115812.

34. R. Yan, C. Gu, D. You, et al., Decoding dynamic epigenetic landscapes in human oocytes using single-cell multi-omics sequencing. Cell Stem Cell, 2021, 28, 1641–1656.e7.

35. Y. Hao, S. Hao, E. Andersen-Nissen, et al., Integrated analysis of multimodal single-cell data. Cell, 2021, 184, 3573–3587.e29.

36. R. Pascual, C. Segura-Morales, M. Omerzu, et al., mRNA spindle localization and mitotic translational regulation by CPEB1 and CPEB4. RNA, 2021, 27, 291–302.

37. J. Huntriss, J. Lu, K. Hemmings, et al. Isolation and expression of the human gametocyte-specific factor 1 gene (GTSF1) in fetal ovary, oocytes, and preimplantation embryos. J. Assist. Reprod. Genet., 2017, 34, 23–31.

38. Y. Zhai, G. Yao, F. Rao, et al., Excessive nerve growth factor impairs bidirectional communication between the oocyte and cumulus cells resulting in reduced oocyte competence. Reprod. Biol. Endocrinol., 2018, 16, 28.

39. D. Bauermeister, M. Claußen, T. Pieler, A novel role for Celf1 in vegetal RNA localization during *Xenopus* oogenesis. Dev. Biol., 2015, 405, 214–224.

40. Y. Wu, H. Fan, Revisiting ZAR proteins: the understudied regulator of female fertility and beyond. Cell. Mol. Life Sci., 2022, 79, 92.

41. J. H. Bergmann, J. Li, M. A. Eckersley-Maslin, et al., Regulation of the ESC transcriptome by nuclear long noncoding RNAs. Genome Res., 2015, 25, 1336–1346.

42. I. Virant-Klun, S. Leicht, C. Hughes, et al., Identification of maturation-specific proteins by single-cell proteomics of human oocytes. Mol. Cell. Proteomics, 2016, 15, 2616–2627.

43. Z. Li, M. Huang, X. Wang, et al., Nanoliter-scale oil-air-droplet chip-based single cell proteomic analysis. Anal. Chem., 2018, 90, 5430–5438.

44. Y. Zhang, Z. Yan, Q. Qin, et al., Transcriptome landscape of human folliculogenesis reveals oocyte and granulosa cell interactions. Mol. Cell, 2018, 72, 1021–1034.e4.

