## Supplementary information for scSTAP analysis for "Simultaneous transcriptome and proteome profiling in a single mouse oocyte with a deep single-cell multi-omics approach"

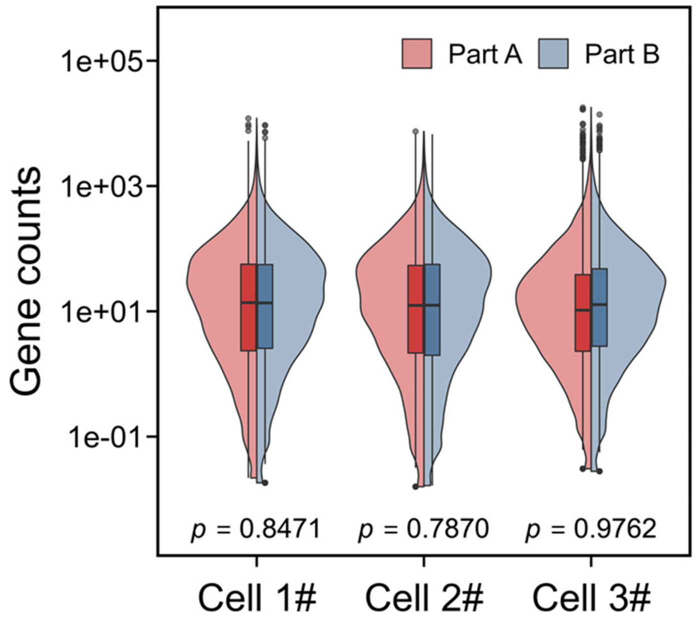


**Figure S1.** Differential expression of genes in the two aliquots of single oocyte samples using single-cell transcriptome analysis with the enzyme-assisted lysis strategy. The results of paired *t*-test showed that there were no significant differences between the genes expression in the two aliquots (n = 3, $p1 = 0.8471$, $p2 = 0.7870$, $p3 = 0.9762$).

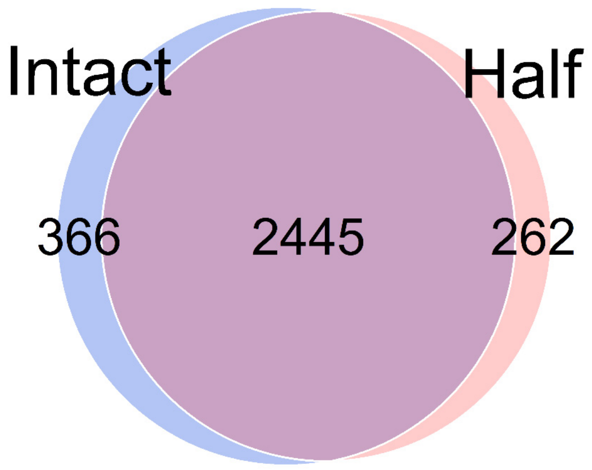


**Figure S2.** Venn diagram of the identified proteins in an intact single oocyte and a half of single oocyte samples. The results showed that there was no significant difference between the identification depth of the whole and the half of cells (n = 3).

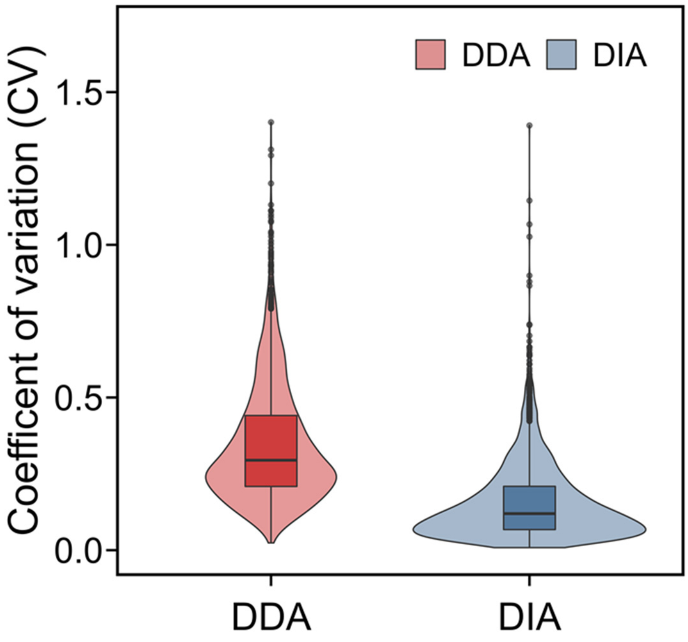


**Figure S3.** Variation coefficient (CV) distribution of the quantified protein groups using the data-dependent acquisition (DDA) and data-independent acquisition (DIA) mode. The median CV of the data under the DDA mode was 29.5% (n = 5), while the median CV of the data under the DIA mode was 12.0% (n = 5).

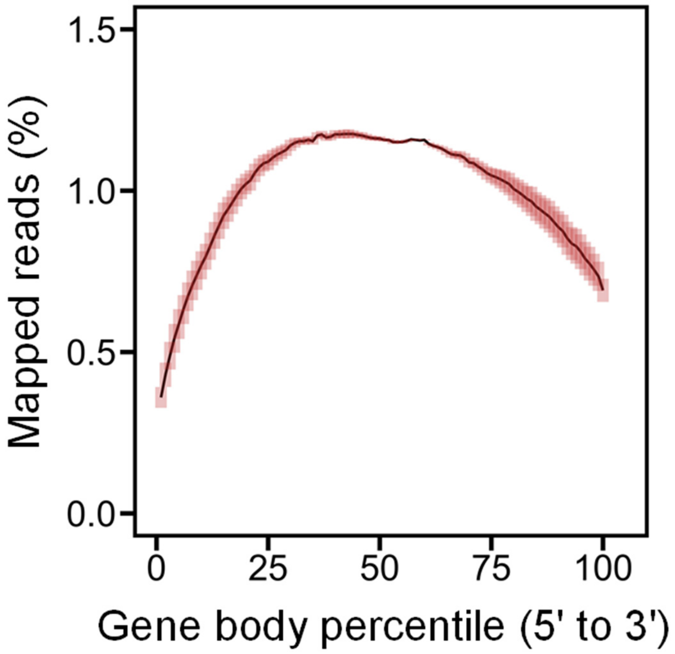


**Figure S4.** Coverage of mapped reads along the gene body for the MATQ-seq method (shaded area: standard deviation of the coverage, oocytes at GV stage, n = 3). There was no obvious 3′- or 5′-end bias observed in the coverage of the mapped reads.

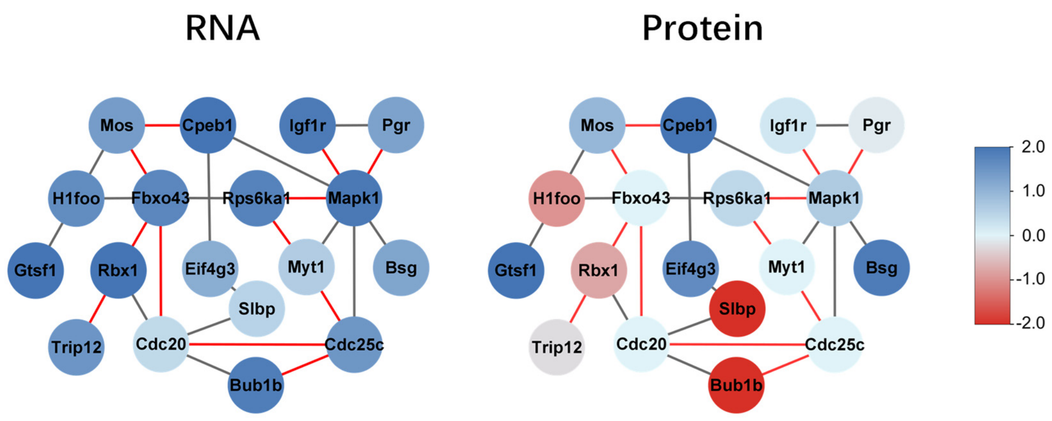


**Figure S5.** Protein-protein interaction analysis of key genes in the "Oocyte Meiosis" pathway with the differential expressions of exons and proteins during oocyte meiotic maturation. Specifically, the red nodes indicated the upregulated genes of single oocytes from GV stage to MII stage, while the blue nodes indicated the downregulated genes during the oocyte meiotic maturation. In addition, the red lines indicated that the regulatory relationship between the two genes were reported in the Kyoto Encyclopedia of Genes and Genomes (KEGG) pathway database.
